# Diet composition drives tissue-specific intensity of murine enteric infections

**DOI:** 10.1101/2023.09.22.558956

**Authors:** Helene Israelsen, Amalie Vedsted-Jakobsen, Ling Zhu, Aurelie Gagnaire, Alexandra von Munchow, Nina Polakovicova, Angela H. Valente, Ali Raza, Audrey I.S. Andersen-Civil, John E. Olsen, Laura J. Myhill, Peter Geldhof, Andrew R. Williams

## Abstract

Diet composition plays a large role in regulating of gut health and enteric infection. In particular, synthetic ‘Western-style’ diets may predispose to disease, whilst whole-grain diets containing high levels of crude fiber are thought to promote gut health. Here we show that, in contrast to this paradigm, mice fed unrefined chow are significantly more susceptible to infection with *Trichuris muris*, a caecum-dwelling nematode, than mice given refined, semi-synthetic diets (SSD). Moreover, mice fed SSD supplemented with inulin, a fermentable fiber, developed chronic *T. muris* burdens whereas mice given SSD efficiently cleared the infection. Diet composition significantly impacted infection-induced changes in the host gut microbiome. Mice infected with the bacterium *Citrobacter rodentium* were also more susceptible to pathogen colonization when fed either chow or inulin-enriched SSD. However, transcriptomic analysis of tissues from mice fed either SSD or inulin-enriched SSD revealed that, in contrast to *T. muris*, increased *C. rodentium* infection appeared to be independent of the host immune response. Accordingly, exogenous treatment with IL-25 partially reduced *T. muris* burdens in inulin-fed mice, whereas IL-22 treatment was unable to restore resistance to *C. rodentium* colonization. Diet-mediated effects on pathogen burden were more pronounced for large intestine-dwelling pathogens, as effects on small intestinal helminth (*Heligmosomoides polygyrus*) were less evident, and protozoan (*Giardia muris*) infection burdens were equivalent in mice fed chow, inulin-enriched SSD, or SSD, despite higher cyst excretion in chow-fed mice. Collectively, our results point to a tissue- and pathogen-restricted effect of dietary fiber levels on enteric infection intensity.

**Importance:** Enteric infections induce dysbiosis and inflammation and are a major public health burden. As the gut environment is strongly shaped by diet, the role of different dietary components in promoting resistance to infection is of interest. Whilst diets rich in fiber or whole grain are normally associated with improved gut health, we show here that these components predispose the host to higher levels of pathogen infection. Thus, our results have significance for interpreting how different dietary interventions may impact on gastrointestinal infections. Moreover, our results may shed light on our understanding of how gut flora and musical immune function is influenced by the food that we eat.

## Introduction

Enteric parasitic and bacterial infections are among the top ten causes of death worldwide. More than 1.5 billion people are infected globally with parasitic worms (helminths), with millions more afflicted with bacterial infections such as *Escherichia coli, Clostridium difficile*, and *Salmonella* (1-3). These pathogens are also ubiquitous in livestock production systems worldwide, threatening food security and economic development (4, 5). Antibiotic and anti-parasitic drug resistance limit effective treatment options for many of these pathogens, and thus novel solutions are urgently required (5, 6).

There is increasing evidence that diet composition plays a key role in mediating resistance to enteric infection and inflammation. Worldwide, there is an increasing shift towards consumption of refined, semi-synthetic diets (SSD), or ‘Western diets’ that lack crude plant fibers and phytochemicals (7). This may have profound consequences for the composition of the gut microbiota, and for the activity of immune cells that reside at the mucosal barrier (8, 9). High rates of obesity and chronic diseases such as colitis may be attributed, in part, to the consumption of refined diets and associated dysbiosis and dysregulated immune function (10, 11).

Nutritional interventions may be a sustainable tool to promote resistance to enteric pathogens through modulation of mucosal immunity. Mucosal immune responses can be broadly classified as type-1, type-2, and type-3; type-1 responses protect against viruses and intracellular bacteria; while type-2 responses defend the host against parasitic worms (helminths); and type-3 responses being crucial against extracellular bacteria and fungi (12, 13). These different arms of the immune response are inter-regulated, and therefore the relative balance of these opposing immune subsets can determine the outcome of infection. The activity of immune cells (e.g. T-cells) in the gut is also continually calibrated by exposure to extrinsic factors such as diet and diet-derived metabolites (14). Dietary components may impact mucosal immune function through several overlapping mechanisms. First, breakdown of dietary carbohydrates and amino acids may favor the growth of defined gut bacteria which subsequently directly impact immune cell behavior (15). Second, altered gut microbiota composition resulting from changing dietary patterns can modulate the abundance of certain gut metabolites such as secondary bile acids, which can impact on immune cell function, e.g. by inducing T-regulatory cells (16). Third, many dietary compounds, e.g. polyphenols or carbohydrate polymers, can directly interact with intestinal epithelial cells or innate immune cells such as dendritic cells to shape the immune and inflammatory tone in the gut (17-19).

Levels of dietary macro- and micro-nutrients such as protein and zinc are known to markedly impact on expression of immunity to enteric infections (20, 21). In addition, a number of studies in both pre-clinical models and human intervention studies have shown proof-of-concept that the gut microbiome and immune function can be substantially modulated by consumption of defined dietary components such as soluble fibers, with potential benefits for metabolic disease and chronic inflammation (22-25). These and other studies have generally defined a paradigm; diets rich in crude fiber and unrefined plant material (e.g. rodent chow) promote a healthy gut and robust and balanced immune function, whilst purified SSD lacking in fiber can lead to dysbiosis, inflammation, and disease, especially when combined with a high level of fat and/or sucrose. However, the interactions between dietary intake and resistance to enteric infection are not yet clear. The composition of dietary fiber has been shown to have a significant influence on the severity of different infections in mice, but sometimes with conflicting results. For example, SSD has been shown to result in more severe infections with the gram-negative bacterium *Citrobacter rodentium* in mice compared with chow, but in other models, infections with *Salmonella* or *Listeria* are enhanced in mice fed chow, relative to mice fed SSD (26, 27). Further studies are clearly required to understand in greater detail the relationships between diet composition, mucosal immunity and enteric infection.

Here, we show that infections in the large intestine with either the helminth parasite *Trichuris muris* or the bacterium *C. rodentium* are substantially lower when mice are fed SSD, compared to either chow or SSD enriched with the soluble fiber inulin. We further demonstrate that whilst increased susceptibility to *T. muris* is associated with altered immune function and can be partially restored with exogenous administration of a type-2 polarizing cytokine, susceptibility to *C. rodentium* occurs despite an intact immune response. Finally, we show that colonization of pathogens residing in the small intestine, rather than in the caecum or colon, are less sensitive to regulation by dietary composition. Taken together, our results shed further light on diet-mediated regulation of responses to mucosal pathogens in the large and small intestinal tissues.

## Results

### Feeding chow increases enteric pathogen burdens relative to semi-synthetic diets

During a long-term project to explore the relationship between diet composition and immunity to helminths, we observed a trend in our laboratory for *T. muris* burdens to be noticeably lower in mice fed experimental open-source SSD, compared to standard rodent chow. To examine this in detail, we utilized a chronic *T. muris* infection model in C57BL/6 mice where animals were infected with a low dose (20 eggs) repeatedly over several weeks, and fed either chow or SSD. Mice were fed SSD or chow for two weeks prior to the commencement of infection, and killed 35 days post-infection (p.i.) after the first egg dose (i.e. 7 weeks total of dietary treatment). This revealed that worm burdens were more than four-fold higher in chow-fed mice, with a substantial number of SSD-fed mice being worm-free (**Figure 1A**). Analysis of earlier time-points showed that at day 11 p.i., all mice fed SSD harbored worms (albeit significantly less so than chow-fed mice), but by day 21 p.i. only very few worms were found in SSD-fed mice, indicating an accelerated expulsion process, rather than reduced establishment (**Supplementary Figure 1**). Furthermore, infected mice fed SSD had significantly lower *T.muris*-specific IgG2a and significantly higher IgG1 antibodies in serum, indicative of a shift from a Th1 to a Th2 response (**Figure 1B**). Regardless of diet, *T. muris* infection significantly increased the expression of *Ifng* and *Il13* in caecal tissue, but the increase in *Ifng* expression was significantly more pronounced (∼1000 fold) in the chow-fed mice than SSD-fed mice (**Figure 1C**). Interestingly, the increase in *Il13* expression was also relatively higher in chow-fed mice than SSD-fed mice, however not to the same as extent as *Ifng,* indicating that the transcriptional response in chow-fed mice was polarized towards a type-1 response (**Figure 1C**). As chronic *T. muris* infection induces crypt hyperplasia (28), we examined the colon for crypt lengths and goblet cell numbers. Infection significantly increased crypt length in mice fed both diets (**Figure 1D**). In accordance with the low worm numbers, crypt lengths were significantly shorter in SSD-fed mice during *T. muris* infection, but also in SSD-fed uninfected mice, indicating that diet composition had a regulatory effect on the gut mucosal tissue independent of infection (*p <* 0.05 for main effects of diet and infection; **Figure 1D**). Consistent with this, goblet cells were lower in uninfected mice fed SSD compared to chow, but not in *T. muris* -infected mice (*p* < 0.05 for interaction between infection and diet). Thus, *T. muris* trickle infection did not increase goblet cell numbers in chow-fed mice, consistent with previous reports (28), but did do so in SSD-fed mice. Furthermore, goblet cell size was also lower in uninfected mice fed SSD, but was boosted by *T. muris* infection, indicating that infection induced an active goblet cell hyperplasia only in mice fed SSD (**Figure 1D**). We noted a similar trend with T-cell populations in mesenteric lymph nodes (MLN). In chow-fed mice, *T. muris* infection did not change the proportion of Th1 (Tbet^+^) cells or Th2 (GATA3^+^) cells (**Figure 1E-F**). In SSD-fed mice, proportions of Th1 cells were significantly lower compared to chow-fed mice in the absence of infection, but were increased in *T. muris-* infected mice (*p*<0.05 for interaction between diet and infection; **Figure 1E**). Th2 cells were not affected by diet in uninfected mice, but were significantly higher in *T. muris-* infected mice fed SSD, relative to infected mice fed chow (*p*<0.05 for interaction between diet and infection; **Figure 1F**), consistent with a type-2 immune bias occurring in SSD-fed mice during *T. muris* infection. Furthermore, we observed that feeding SSD significantly decreased Treg (Foxp3^+^) populations independently of *T. muris* infection (*p*<0.05 for main effect of diet; **Figure 1F**). Notably, these effects of diet on T-cell populations appeared to result from prolonged consumption of the diets, as at the time point trickle infection was commenced (i.e. after 14 days diet adaptation), no differences were observed in T-cell subsets (**Supplementary Figure 2**). Collectively, these data show that diet composition substantially impacted *T. muris* infection levels, and that indicators of an activated type-2 immune response were more evident in SSD-fed mice than in chow-fed mice.

**Figure 1.**
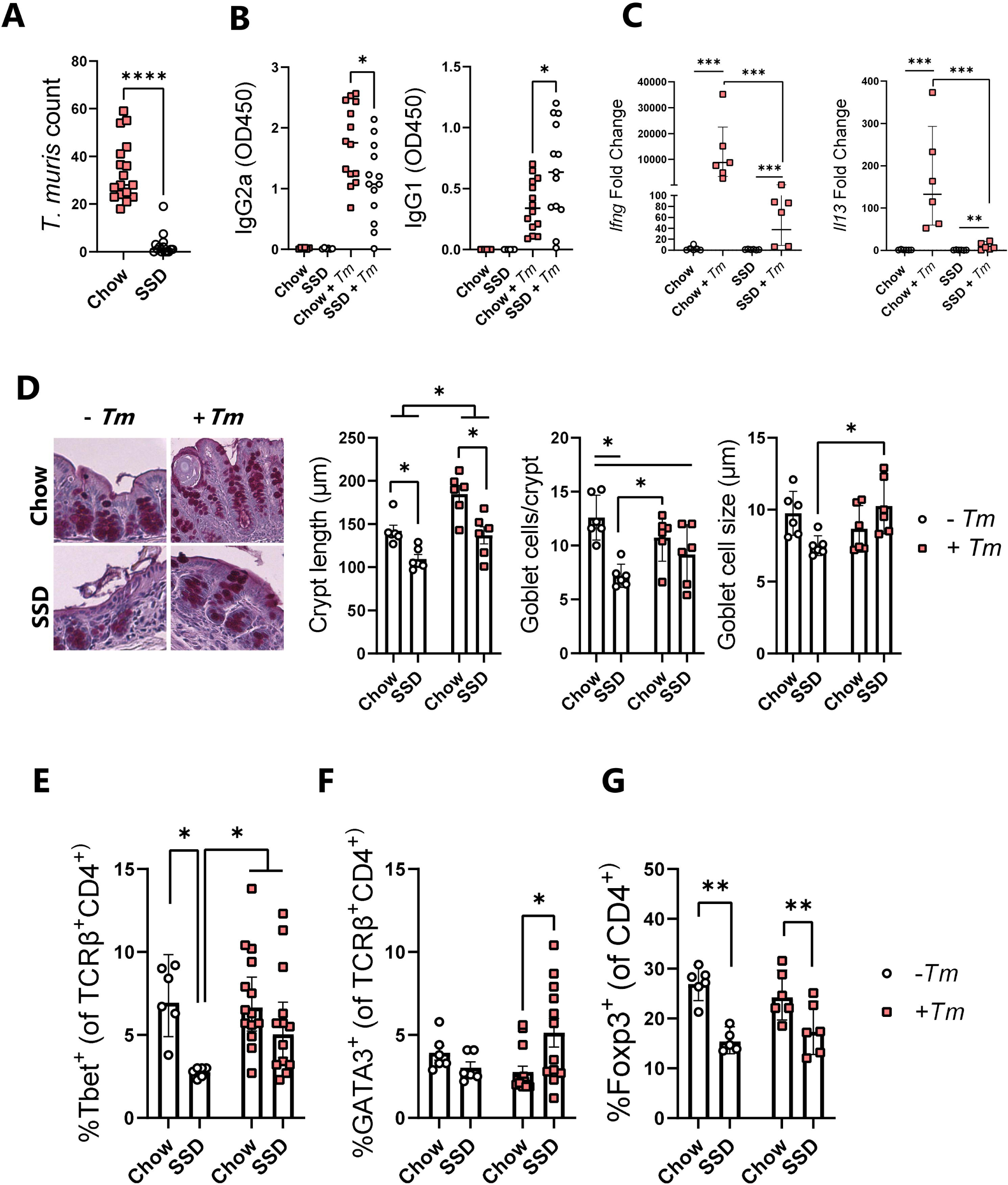
Impact of chow or semi-synthetic diets on *Trichuris muris* infection burdens and host immune response. Mice were fed either chow or semi-synthetic diets (SSD), trickle-infected with *T. muris* by inoculation with 20 eggs on day 0, 7 and 14, and were sacrificed on day 35 post-infection (p.i.). **A)** Worm burdens in caecum on day 35 p.i. by. Data is from three independent experiments, each with *n*=3-8 mice per dietary group. **B)** *T. muris*-specific IgG2a and IgG1 levels in serum on day 35 p.i. Data from infected mice is from two independent experiments, each with 6-8 mice per treatment group, and data from uninfected mice is from a single experiment with n=6 mice per group. **C)** Expression of *Ifng* and *Il13* assessed by qPCR in the caecum day 35 p.i. Data are from a single experiment, with n=6 mice per treatment group. Fold changes are relative to uninfected controls within each diet group. **D)** Crypt hyperplasia, goblet cell numbers and goblet cell diameter in colon samples at day 35 p.i. Data are from a single experiment, with n=6 mice per treatment group Percentage of Tbet^+^ (**E**) and GATA3^+^ CD4^+^ T-cells (**F**), and Foxp3^+^ CD4^+^ cells (**G**), in the mesenteric lymph nodes of mice on day 35 p.i. For Tbet^+^ and GATA3^+^ expression, data from infected mice is from two independent experiments, each with 5-8 mice per treatment group, and data from uninfected mice is from a single experiment with n=6 mice per group. For Foxp3^+^ expression, data are from a single experiment, with n=5-6 mice per treatment group. See **Supplementary Figure 4** for T-cell gating strategy. **** *p* < 0.001, \**p* < 0.05 by Mann-Whitney test, comparing infected groups on chow and SSD (**A**,**B**). \*\*\**p*<0.005, \*\**p*<0.01, *p*<0.05 by two-way ANOVA, with Tukey’s post-hoc testing where interaction term is significant (**C-G**). Data are presented as medians (**A,B**), geometric means with 95% confidence intervals (**C, E**) or mean ± S.E.M. (**D, F-G**).

Due to our observation that the increased *T. muris* burdens in chow-fed mice were associated with reduced type-2 immune function, we speculated that SSD may enhance type-1 or type-3/17 responses due to the reciprocal regulation of T-helper cell subsets (29), and thus promote resistance to a pathogen where these responses are important. Contrary to our expectations, mice infected with the bacterial pathogen *C. rodentium,* where immunity is dependent on type-3-driven induction of IL-22 and expression of anti-microbial peptides such as Reg3γ, also had significantly higher pathogen burdens when fed chow compared to SSD (**Figure 2A**). This effect was also associated with increased levels of *C. rodentium*-specific IgG2a in serum (**Figure 2B**). However, expression of *Il22* and *Nos2* (encoding the inducible nitric oxide synthase) in the colon, both genes known to be strongly induced by *C. rodentium* infection (30, 31) was equivalent between dietary treatments (**Figure 2C-D**). Thus, SSD consumption was associated with reduced susceptibility to two different enteric pathogens.

**Figure 2.**
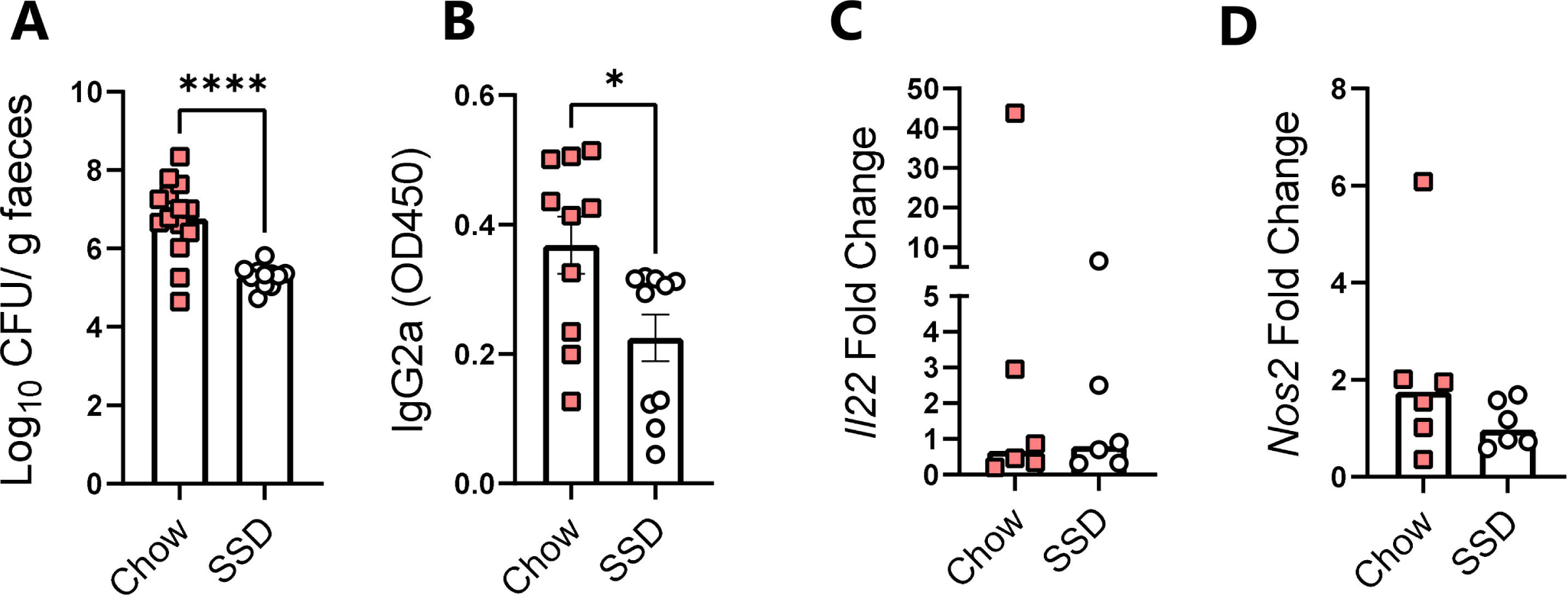
Impact of chow or semi-synthetic diets on *Citrobacter rodentium* infection burdens and host immune response. Mice were fed either chow or semi-synthetic diets (SSD), infected with *C. rodentium* and sacrificed 8 days post-infection (p.i.). **A)** Faecal *C. rodentium* burdens on day 8 p.i. **** *p* < 0.001 by unpaired t-test. **B)** *C. rodentium*-specific IgG2a levels in serum on day 8 p.i. * *p* < 0.05 by unpaired t-test. **C-D)** Expression of *Il22* and *Nos2* in colon on day 8 p.i. Fold changes are relative to the chow-fed infected mice. Data is from three independent experiments, each with *n*=3-6 mice per dietary group (**A**), two independent experiments, each with 6 mice per group (B) or a single experiment with *n*=6 per dietary group (**C-D**), and presented as mean ± S.E.M. (**A-**B) or medians (**C-D**).

### Enriching semi-synthetic diets with inulin increases pathogen burdens

Chow is a complex mixture of different components with a high level of crude, soluble fibers, in contrast to SSD which is devoid of soluble fiber and instead contains only insoluble cellulose. We therefore examined the effect of enriching SSD with inulin, a highly soluble fiber, on infection levels. We have previously found that including inulin in SSD increased *T. muris* burdens in mice during acute infection (a single, high dose of 300 eggs), whereas cellulose fiber had no effect (32). To explore if trickle-infected mice also had higher *T. muris* burdens when fed inulin, mice were fed either SSD or inulin-enriched SSD and given the same trickle infection regime as above. Strikingly, we observed that mice fed inulin also had a substantially higher burden of *T. muris* than mice fed SSD alone (**Figure 3A**), with the effect being very similar to that of chow. Furthermore, in line with chow experiments, *C. rodentium* mice fed inulin-enriched SSD had higher infection burdens than those fed SSD (**Figure 3B**), which is consistent with a recent report (33). Thus, feeding either unrefined chow, or SSD with an added source of purified soluble fiber, was sufficient to increase infection with two evolutionary divergent enteric pathogens.

**Figure 3.**
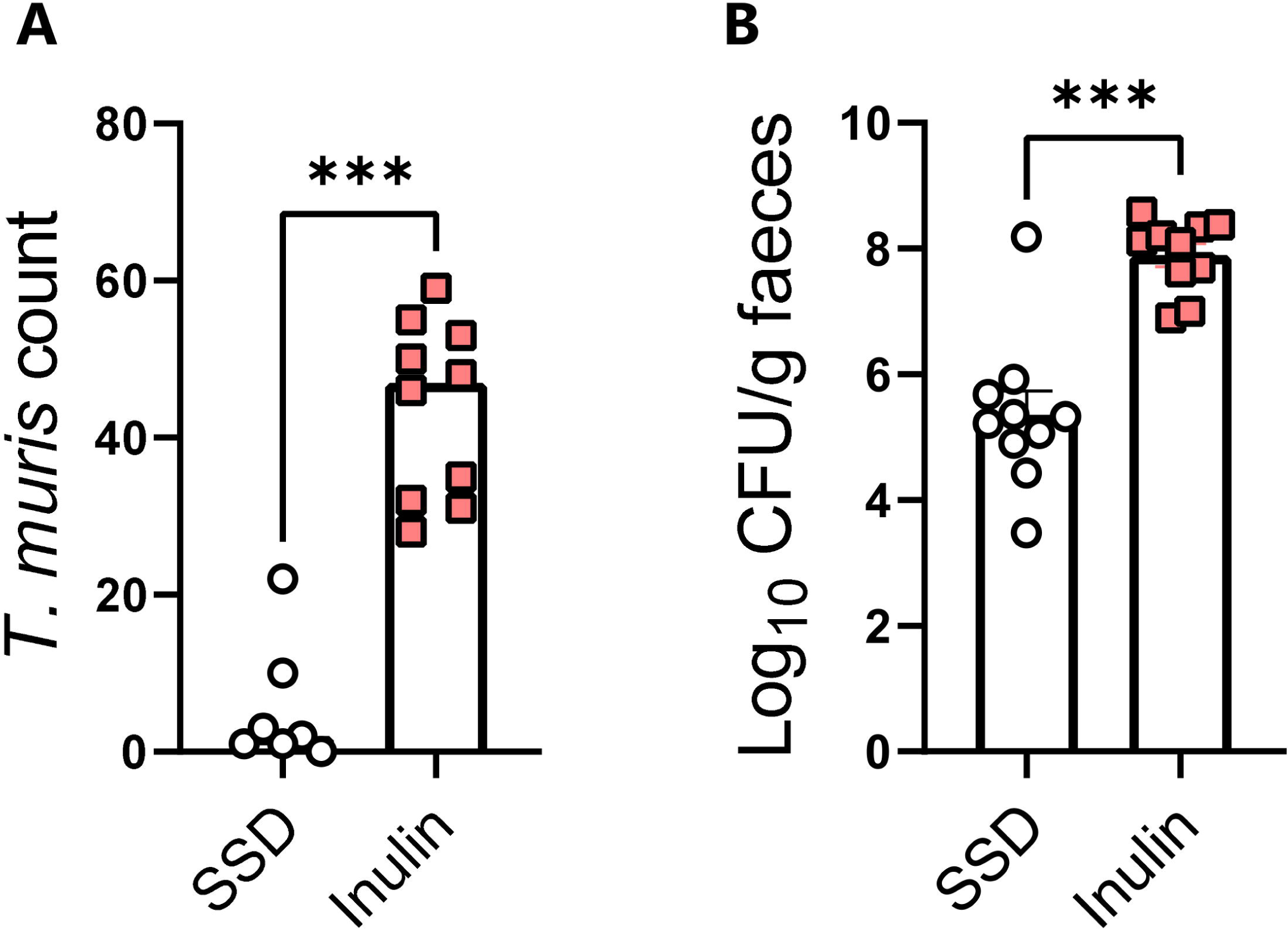
Impact of semi-synthetic diets with or without inulin-enrichment on *Trichuris muris* and *Citrobacter rodentium* burdens. Mice were fed either semi-synthetic diets alone (SSD), or SSD enriched with 10% inulin (Inulin). Mice were trickle-infected with *T. muris* by inoculation with 20 eggs on day 0, 7 and 14, and sacrificed on day 35, or infected with *C. rodentium* and sacrificed on day 8 post infection (p.i.). **A)** Caecal *T. muris* burdens on day 35 p.i. Data are from two independent experiments with *n*=3-6 mice per treatment group, and shown as median values. *** *p* < 0.001 by Mann-Whitney test. **B)** Faecal *C. rodentium* burdens at day 8 p.i. *** *p* < 0.001 by unpaired t-test. Data are from two independent experiments, each with *n*=3-6 mice per treatment group,, and shown is mean ± S.E.M.

### Diet composition restructures the gut microbiome during Citrobacter rodentium and Trichuris muris infection

To further explore the effect of the different diets on the gut environment, faecal samples taken from mice fed chow, SSD, or inulin-enriched SSD with either *C. rodentium* or *T. muris* infection, and their respective uninfected controls, were analyzed for gut microbiome (GM) composition by 16S rRNA amplicon sequencing. Strong interactions were observed between diet and infection. α-diversity tended to be decreased by *C. rodentium* infection, at both observed species and Shannon index level (main effect of infection, *p* = 0.07 and *p* = 0.052, respectively, **Figure 4A**). Diet had a substantially stronger effect with α-diversity metrics significantly higher in chow-fed mice than those fed SSD, which likewise had a more diverse GM than inulin-fed mice (*p*<0.0001 for main effect of diet; **Figure 4A**). β-diversity also differed significantly as a result of both diet and infection (*p*<0.05 for interaction between diet and infection, PERMANOVA on Bray-Curtis distance metrics; **Figure 4B**). Pairwise testing revealed that infection tended (adjusted *p* = 0.07) to affect the GM only in chow-fed mice, and not in SSD-fed or inulin-fed mice. Diet had a strong effect on the GM composition which was largely independent of infection. In uninfected mice, the composition in SSD-fed animals was significantly different from those fed both chow or inulin (adjusted *p*<0.05), and chow and inulin also tended to result in distinct compositions (adjusted *p*=0.051). Within infected mice, the same trend was evident but only significantly so for chow and SSD (adjusted *p* values <0.05). At the genus level, the main compositional changes resulting from diet were a substantial reduction in the relative abundance of *Lactobacillus* and *Muribaculaceae* in mice fed SSD (with or without inulin enrichment) compared to chow, and a corresponding increase in *Faecalbacterium* and *Bifidobacterium* (**Figure 4C-D**). Within the SSD diets, inulin enrichment resulted in an increase in *Faecalbacterium* and *Muribaculum* and a reduction in *Clostridum senso stricto 1.* These changes were further impacted by infection. For example, infection tended to increase *Lactobacillus* only in chow-fed mice, and *Bifidobacterium* only in inulin-fed mice, whilst reductions in *Muribaculum* and *Clostridum senso stricto 1* were apparent only in chow-fed mice (**Figure 4C-D**). Overall, these data demonstrate that the different diets had a substantial effect on the underlying GM composition, and that they influenced the response to *C. rodentium* infection, with only the GM of chow-fed and inulin-fed mice actively changing as a response to the pathogen.

**Figure 4.**
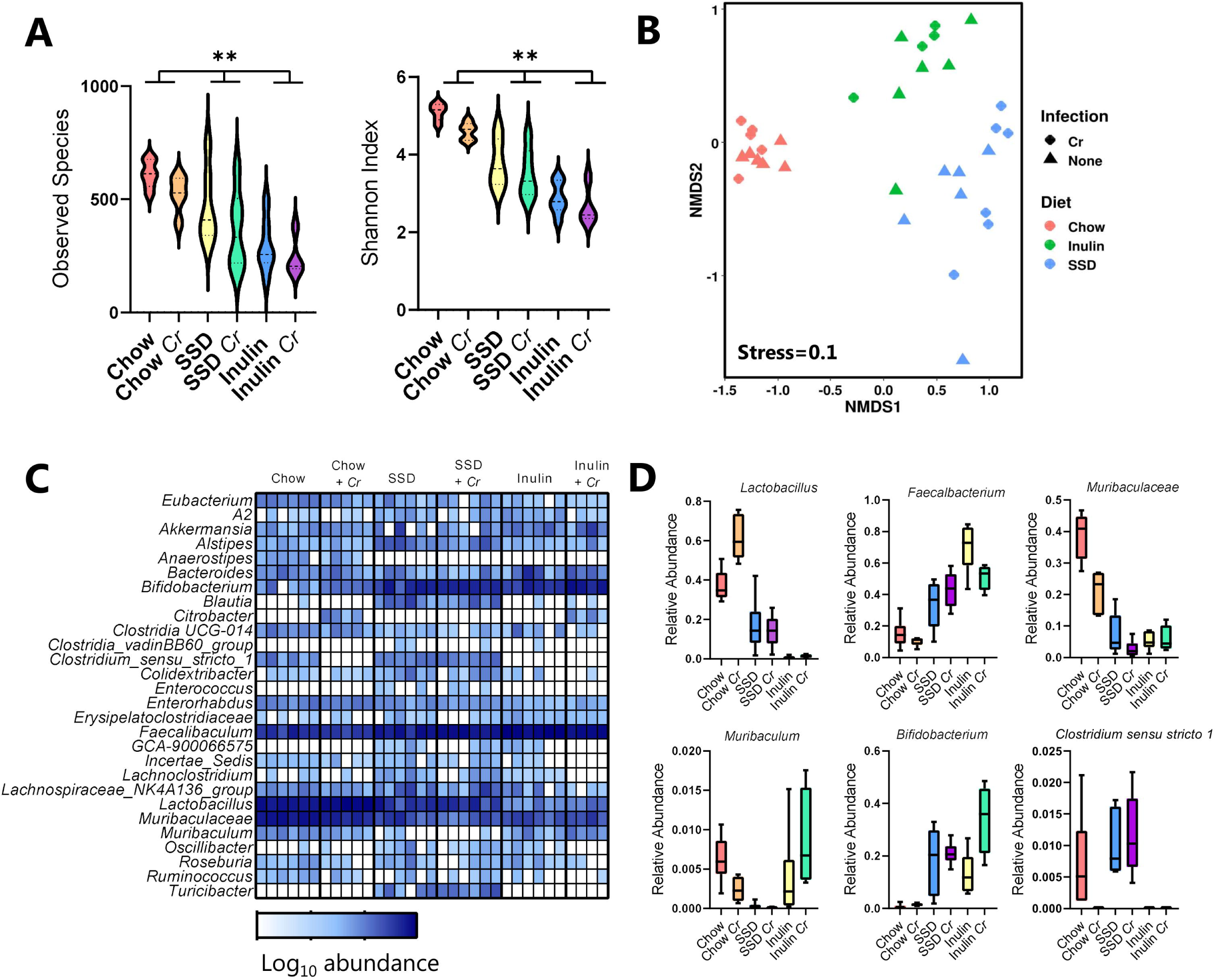
Impact of diet and *Citrobacter rodentium* infection on the host gut microbiome. Mice were fed either chow, semi-synthetic diets alone (SSD), or SSD enriched with 10% inulin (Inulin). Mice were infected with *C. rodentium*, or left as uninfected controls, and faecal samples collected at day 7 post-infection. n=4-6 per treatment group. **A)** α-diversity (Observed species and Shannon index), in mice fed chow, SSD, or inulin, with or without *C. rodentium* (*Cr*) infection. ** *p*<0.01 by two-way ANOVA. **B)** Non-metric multidimensional scaling (NMDS) plot showing β-diversity of different treatment groups. Relative abundance at genus level (**C**), and selected genera (**D**) in mice fed chow (C), SSD, or inulin (I), with or without *Cr* infection.

A similar pattern was observed in the *T. muris* experiments. Diet had a strong influence on α-diversity, with uninfected, chow-fed mice clearly having the most diverse GM. Infection significantly decreased this diversity in chow-fed mice, however this effect was not evident in SSD- and inulin-fed mice (*p*<0.05 for interaction between diet and infection; **Figure 5A**). Diet and infection also exhibited strong interactions on β-diversity (*p*<0.05 for interaction between diet and infection, PERMANOVA on Bray-Curtis distance metrics; **Figure 5B**). Diet significantly impacted the GM composition, regardless of infection status (adjusted *p* <0.05 for all PERMANOVA pairwise comparisons between diet groups in both uninfected and *T. muris* -infected mice). In contrast, the effect of infection was diet-dependent, with *T. muris* infection significantly impacting the GM only in chow-fed and inulin-fed mice (adjusted *p* <0.05 for PERMANOVA pairwise comparisons) but not in SSD-fed mice (**Figure 5B**). In the absence of infection, chow diets favored the growth of *Lactobacillus* and *Muribaculaeceae*, whilst SSD (with and without inulin) promoted the growth of *Faecalbacterium*, consistent with the *C. rodentium* study. Notably, *T. muris* infection also promoted *Lactobacillus* abundance that was most pronounced in inulin-fed mice, as well as the relative abundance of *Bifidobacterium* (**Figure 5C-D**). However, the relative abundance of *Entererococcus* and *Escherichia/Shiga* was also increased by *T. muris* infection, most drastically in the inulin-fed mice (**Figure 5C-D**). Furthermore, we noted that infection also significantly decreased the abundance of *Alstipes* spp., most prominently in chow-fed and inulin-fed mice. Thus, consistent with the effects of diet on worm burden, chow and inulin diet most strongly enabled *T. muris*-induced dysbiosis whilst the effect of infection on the SSD diet was far more subtle.

**Figure 5.**
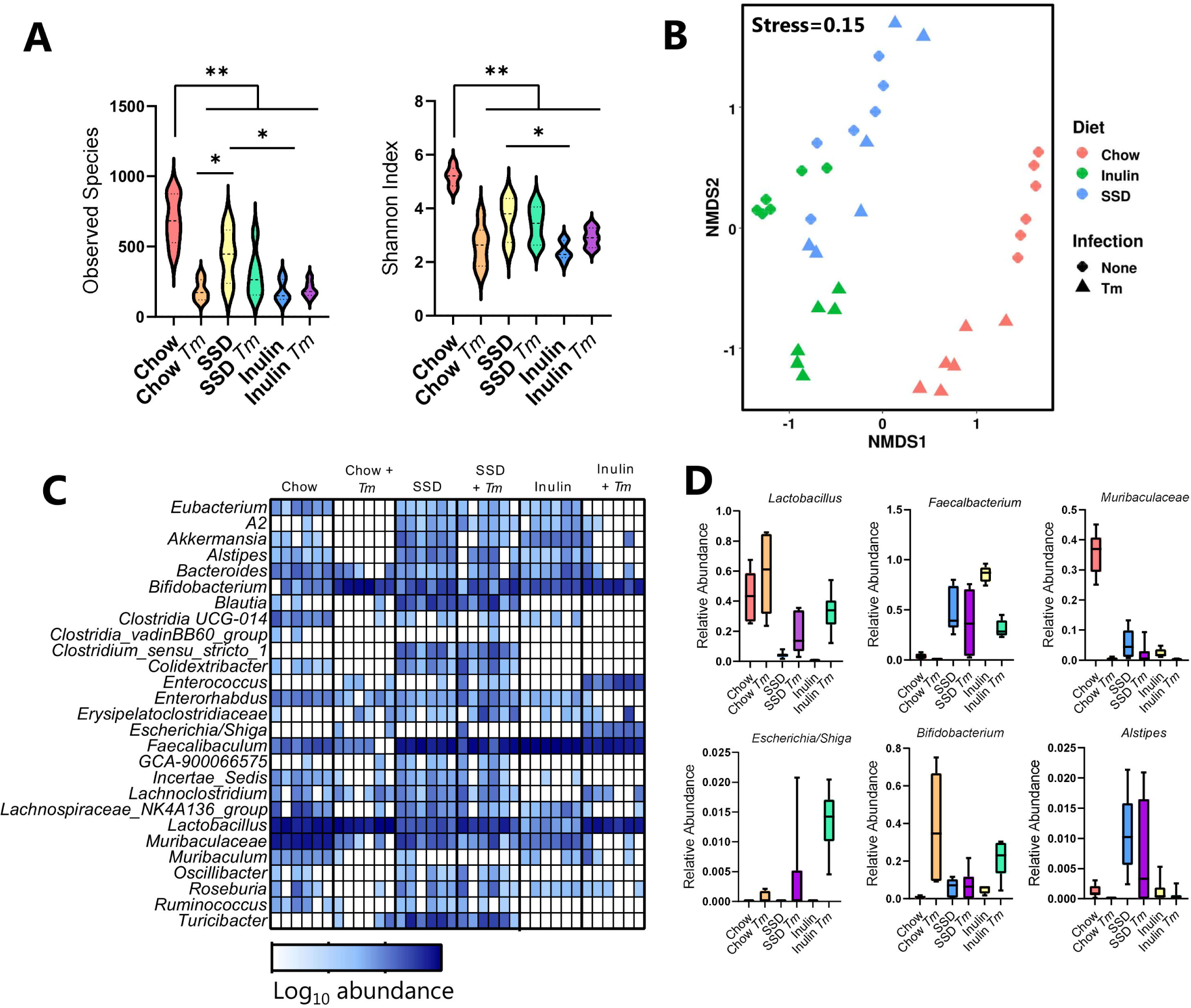
Impact of diet and *Trichuris muris* infection on the host gut microbiome. Mice were fed either chow, semi-synthetic diets alone (SSD), or SSD enriched with 10% inulin (Inulin). Mice were infected with *T. muris*, or left as uninfected controls, and faecal samples collected at day 35 following the commencement of trickle infection (20 eggs given at day 0, 7 and 14 days post-infection). n=6 per treatment group. **A)** α-diversity (Observed species and Shannon index), in mice fed chow, SSD, or inulin, with or without *T. muris* (*Tm*) infection. \*\**p*<0.01, \**p*<0.05 by two-way ANOVA, with Tukey’s post-hoc testing where interaction term is significant. **B)** Non-metric multidimensional scaling (NMDS) plot showing β-diversity of different treatment groups. Relative abundance at genus level (**C**), and selected genera (**D**) in mice fed chow (C), SSD, or inulin (I), with or without *Tm* infection.

### Dietary inulin increases Trichuris muris and Citrobacter rodentium burdens through distinct mechanisms

Given that addition of inulin to SSD was sufficient to increase both *T. muris* and *C. rodentium* burdens, we next sought to determine whether this resulted from a common mechanism. Inulin-mediated susceptibility to acute, high-dose *T. muris* infection can be reversed by neutralization of IFNγ, thus manipulating the type-1/type-2 immune balance in favor of type-2 immunity (32). In contrast, it has recently been reported that higher burdens of *C. rodentium* in mice fed inulin-enriched SSD appear to be independent of the immune response, as similar effects were observed in mice deficient in IL-18, IL-22 or adaptive immune cells (Rag1^-/-^) (33). This suggests that the increased burdens of *T. muris* and *C. rodentium* may reflect two distinct mechanisms. To examine this issue in more detail, we harvested caecal or colon tissue from *T. muris* or *C. rodentium* -infected mice respectively after feeding them with either SSD or inulin-enriched SSD. Transcriptomic profiling of caecum tissue from *T. muris* -infected mice revealed a dramatic remodeling of the host immune response. More than 1900 genes were significantly regulated (adjusted *p* value <0.05, fold change > 2; **Supplementary File**), with many of the upregulated genes being involved in pro-inflammatory or type-1 related immune responses (**Figure 6A**). These included genes such as *Ifng*, *Il27*, and *Cxcl10*, which have all been implicated in inhibiting resistance to *T. muris* (34-36). In contrast, type-2 related genes such as *I L 4*, *IL13* and *Ang4*, all of which have been shown to be important for *T. muris* expulsion (36, 37) were suppressed (**Figure 6A**). Amongst the top ten upregulated genes in inulin-fed mice were *Nos2* and *Gzmk*, encoding a granzyme normally found in cytotoxic T-cells and involved in antibacterial immune responses (**Figure 6B**). Downregulated genes included *Mxpt1,* encoding a mucosal pentraxin, as well as genes encoding other mucosal defense molecules such as intelectins that are important for anti-helminth immunity (**Figure 6B**). Finally, gene-set enrichment analysis (GSEA) revealed that the upregulated gene pathways were related to both cell cycle processes (e.g. DNA Replication), but also cytokine signaling, anti-viral (RIG-I-like receptor signaling) responses, and anti-bacterial (TLR and NOD-like receptor signaling) responses (**Figure 6C**). Downregulated pathways were associated with diverse physiological roles such as calcium signaling and neuroactive ligand receptor interactions (**Figure 6C**). Taken together, these data suggest that inulin promotes an inappropriate type-1-biased immune response during a low-dose trickle infection, which results in parasite chronicity and associated inflammation.

**Figure 6.**
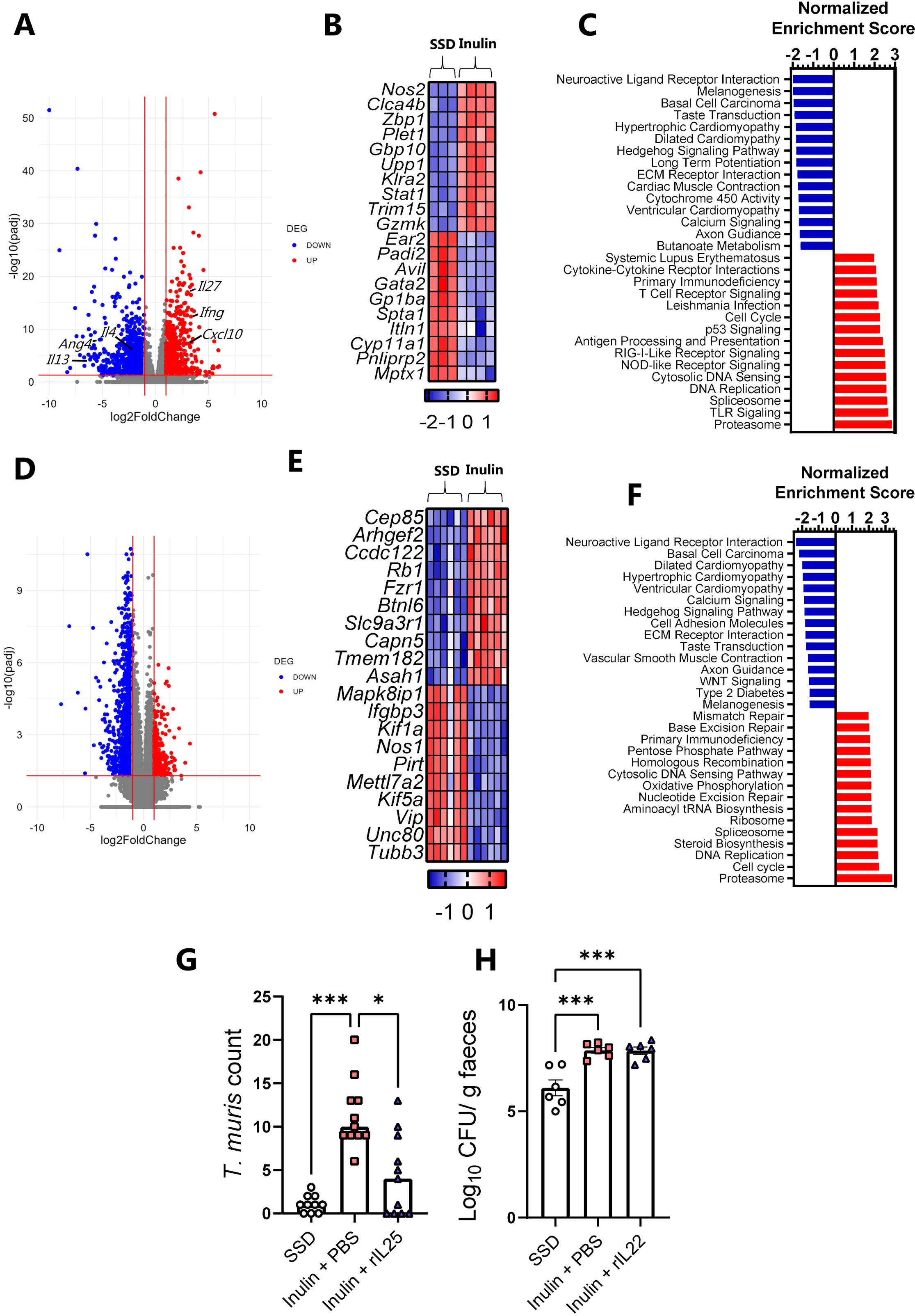
Transcriptomic analysis and cytokine administration reveal independent mechanisms for increased *Trichuris muris* and *Citrobacter rodentium* burdens by dietary inulin. **A-F)** Mice were fed either semi-synthetic diets alone (SSD), or SSD enriched with 10% inulin (Inulin). Mice were trickle-infected with *T. muris* by inoculation with 20 eggs on day 0, 7 and 14, and were sacrificed on day 35 (*n*=3-4 per dietary group; **A-C**), or infected with *C. rodentium* and sacrificed on day 8 post infection (p.i.) (*n*= 6 per dietary group; **D–F**). **A, D)** Volcano plots showing differentially expressed genes (adjusted *p* value < 0.05; Fold change > 2). **B, E)** Heat map showing top ten up- and downregulated genes as assessed by adjusted *p* values. A full list of significantly regulates genes is shown in Supplementary File 1. **C, F)** Top fifteen KEGG pathways enriched or suppressed by feeding inulin relative to SSD (*q* value <0.05), identified by gene-set enrichment analysis. **G)** Worm burdens in caecum on day 21 p.i. following infection with 20 *T. muris* eggs in mice fed SSD, Inulin + vehicle control (PBS), or Inulin + rIL-25. Data are from two independent experiments, each with *n*=5-6 mice per treatment group, and shown as median values. \*\*\**p*<0.005, \**p* < 0.05 by Kruskal-Wallis test followed by Dunn’s post-hoc testing. **H)** Faecal *C. rodentium* burdens on day 8 p.i. in mice fed SSD, Inulin + vehicle control (PBS), or Inulin + rIL-22. *n*=6 mice per treatment group from a single experiment. Data shown as mean ± S.E.M. \*\*\**p*<0.005 by Tukey’s post-hoc testing following one-way ANOVA.

Next, the colonic transcriptomic response to dietary inulin in *C. rodentium* -infected mice was investigated. Relative to mice fed SSD alone, mice fed inulin-enriched SSD had altered expression of approximately 2000 genes (**Figure 6D; Supplementary File**). However, unlike *T. muris*, few of these genes were involved in immune function. Inspection of the top ten up- and downregulated genes showed that most were associated with metabolic and cell cycle processes such as centrosome function (*Cep85*), Rho GTPases (*Arhgef2*), tubulin activity (*Tubb3*) and neuropeptide signaling (*Vip*) (**Figure 6E**). Similar to experiments with *T. muris-* infected mice, downregulated KEGG pathways identified by GSEA included calcium signaling and neuroactive ligand-receptor interactions. A number of pathways related to cardiac myopathy were also inhibited by inulin in both infection models, as a consequence of the suppression of many genes encoding voltage-dependent calcium channels (e.g. *Cacng4*) and cAMP formation (e.g. *Adcy3*). Enriched gene pathways included proteasome and spliceosome function (both also observed in *T. muris* -infected mice) as well as pathways related to ribosome activity and nucleotide excision and repair (**Figure 6F**). A closer inspection of genes known to be involved in immunity to *C. rodentium* (38, 39) revealed no significant suppression related to inulin intake (**Supplementary File**). For instance, *Il22* and *Reg3g* expression was not affected by inulin, whilst expression of *Il18* was significantly enhanced (fold change +1.5), and *Il27* also tended to be enhanced (fold change +2.6; adjusted *p* value 0.13). Genes encoding beta-defensins were either unchanged or in the case of *Defb45* were significantly enhanced in inulin-fed mice (data not shown). Overall, these results indicate that whilst inulin-mediated enhancement of *T. muris* infection was associated with impaired immune function, enhancement of *C. rodentium* infection was manifested despite a seemingly intact immune response, suggesting that the mechanisms through which diet increases pathogen burdens are different.

To test this in more detail, we reasoned that strengthening the type-2 response in *T. muris*-infected mice fed inulin should restore resistance to infection, whereas equivalent support of the immune response in inulin-fed *C. rodentium*-infected mice would not have the same effect as an appropriate immune response was already generated. To this end, we fed mice inulin-enriched SSD and gave them a single dose of 20 *T. muris* eggs. Mice were then treated with exogenous IL-25, a cytokine which stimulates a broad type-2 immune response in mucosal tissues (40), or vehicle control. Compared to mice fed SSD alone, inulin again resulted in a drastic increase in worm burden, whereas mice treated with IL-25 had significantly less worms than inulin-fed mice given vehicle control (**Figure 6G**). Thus, administration of a type-2 polarizing factor was sufficient to lower infection during inulin consumption. In contrast, administration of IL-22 was not able to lower burdens of *C. rodentium* in inulin-fed mice. Mice fed inulin had higher faecal burdens of *C. rodentium* than mice fed SSD (**Figure 6H**), and these burdens were equivalent in IL-22-treated or vehicle control mice. Thus, these data support the notion that dietary inulin impairs type-2 immunity towards *T. muris* infection but enhances *C. rodentium* burdens by a mechanism independent of immune function.

### Feeding chow or inulin-enriched semi-synthetic diets has less pronounced effects on the course of small intestinal pathogen infections

Given that *T. muris* and *C. rodentium* share a relative similar tissue niche in the large intestine, we were also interested to see if these dietary effects were applicable to pathogens residing in the more proximal part of the gut, i.e. the duodenum and jejunum. We selected the helminth *Heligmosomoides polygyrus*and the protozoa *Giardia muris* as two taxonomically distinct pathogens to assess this possibility. We observed that *H. polygyrus* burdens were similar between dietary groups at day 14 p.i., but at day 28 p.i. SSD-fed mice had lower worm burdens than chow-fed mice (but not inulin-fed mice; **Figure 7A**). Faecal egg counts were also lower in SSD-fed mice at day 28 p.i., but not significantly (**Figure 7B**). Assessment of jejunum histology samples at day 28 p.i. showed that ratios of villous height to crypt depth (VCR) were suppressed by *H. polygyrus* infection in chow-fed and inulin-fed mice, but not SSD-fed mice (*p*<0.05 for interaction between diet and infection; **Figure 7C-D**). However, goblet cell numbers and size were increased by *H. polygyrus* comparably in all groups (*p* < 0.05 for main effect of infection, **Figure 7C-D**). Moreover, serum IgG1 levels were equivalent in infected mice fed the different diets (*p* < 0.05 for main effect of infection **Figure 7D**). None of these parameters were affected by diet in the absence of infection (**Figure 7D**). *G. muris* trophozoite counts were equivalent in the small intestine of mice fed either chow, SSD, or inulin-enriched SSD (**Figure 7E**). Interestingly, cyst output was significantly lower in mice fed SSD (with or without inulin), compared to chow (**Figure 7F**). Thus, whilst diet composition also had effects on *H. polygyrus* infection and may influence the transmissibility of small intestinal protozoa, the effects on infection intensity were noticeably lower than with the caecum- and colon-dwelling pathogens,. Whilst we cannot rule out that these differential effects are intrinsic to the different pathogens studied and not their location in the gut, it does raise the possibility that tissue niche has an influence on the regulation of enteric pathogens by dietary factors.

**Figure 7.**
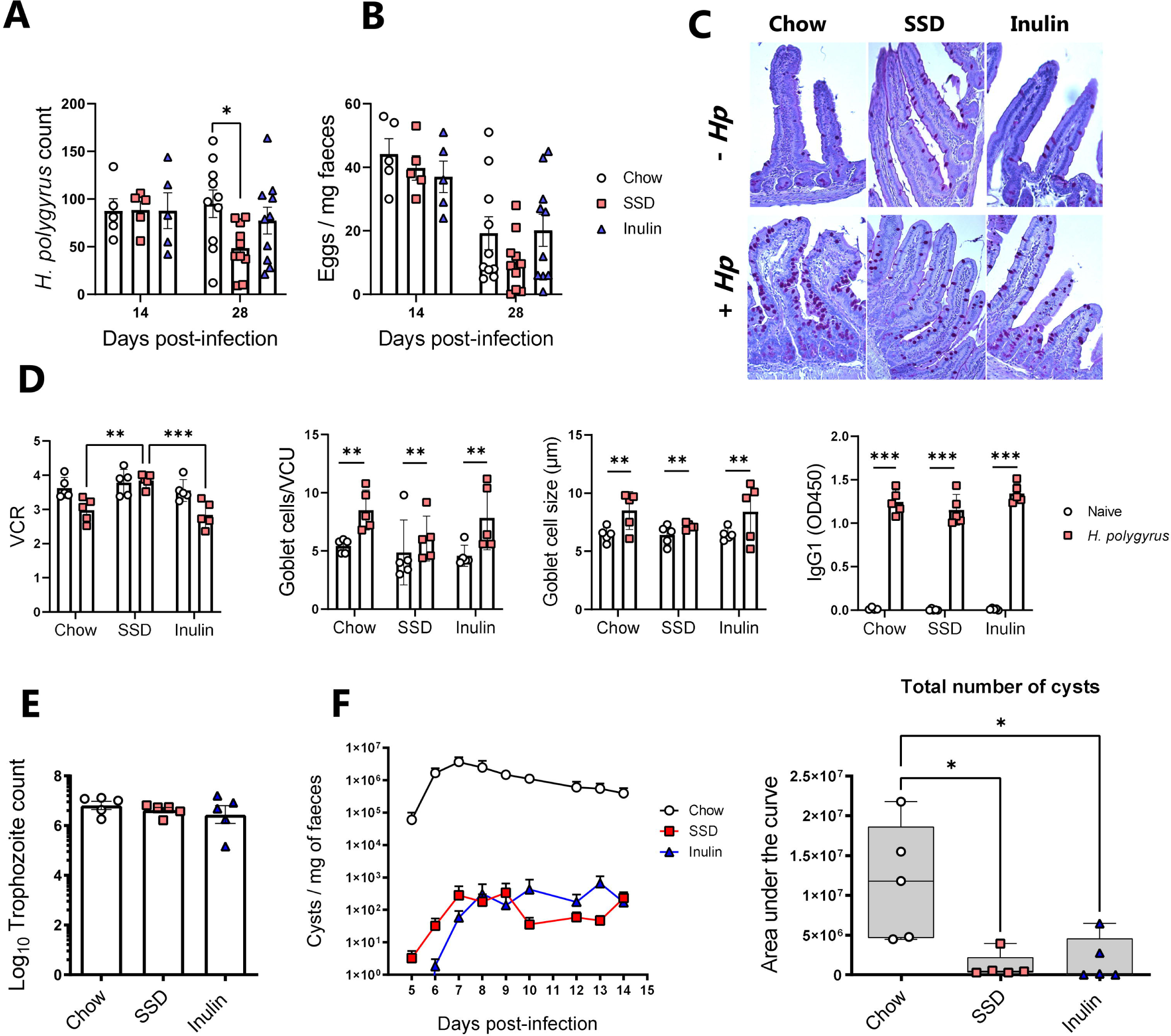
Impact of chow or semi-synthetic diets with or without inulin-enrichment on *Heligmosomoides polygyrus* and *Giardia muris* burdens. Mice were fed either chow, semi-synthetic diets alone (SSD), or SSD enriched with 10% inulin (Inulin). Mice were infected with *H. polygyrus* and sacrificed on either day 14 or day 28 post-infection (p.i.), or *G. muris* and sacrificed on day 16 p.i., Data shown as mean ± S.E.M. **A)** *H. polygyrus* worm burdens in small intestine and **B)** faecal egg counts on day 14 and 28 p.i. \**p*<0.05 by Tukey’s post-hoc testing following one-way ANOVA at each time-point. Data is from two independent experiments, each with n=5 per treatment group (day 28 p.i.), or a single experiment with n=5 (day 14 p.i.) **C-D)** Assessment of villous height to crypt depth ratio (VCR), goblet cell numbers, goblet cell diameter and serum *H. polygyrus* -specific IgG1 levels in the proximal jejunum of mice (day 28 p.i.) with or without *H. polygyrus* infection, and fed different diets. Data is from a single experiment with n=5 per treatment group. \*\*\**p*<0.005, \*\**p*<0.01 by two-way ANOVA, with Tukey’s post-hoc testing where interaction term is significant. **E)** *G. muris* trophozoites counts in the small intestine on day 16 p.i **F)** Daily *G. muris* cyst output in faeces and cumulative cyst counts (total cyst number) over the course of infection from day 5 until day 14 p.i. \**p* < 0.05 by Kruskal-Wallis test and Dunn’s post-hoc testing. Data is from a single experiment with n=5 per treatment group.

### Diet composition regulates the gut microbiota composition and primes mucosal tissue differentially in the colon and small intestine

To further explore whether the different tissue sites (small vs. large intestine) may respond differently to the change in diet composition, we first assessed the mucosal tissue architecture following 14 days of diet adaption (i.e. the point of pathogen infection). In both the jejunum and colon, goblet cell numbers were not affected by the different diets, and VCR in the jejunum were equivalent between dietary groups (**Figure 8A**). However, we noted that crypt lengths in the colon tended to be shorter in mice fed SSD than those fed chow on inulin (*p* = 0.07; **Figure 8A**). Given that we also observed the same phenomenon following extended consumption of SSD during *T. muris* experiments (Figure 1), we speculated that the colon may be a more responsive site to the changes in diet composition. The large intestine (LI) represents the main site of microbial fermentation in the gut, and therefore it may be plausible that increased microbial loads in different tissue sites may differentially affect how these tissues respond to pathogen infection. Therefore, we assessed GM composition in both the small intestine (SI) and LI (caecum and proximal colon) in mice fed chow, SSD, or inulin, for 14 days. α-diversity was lower in mice fed SSD compared to chow, and was also substantially lower in the SI than in the colon, suggesting a richer GM composition in the LI (**Figure 8B**). Moreover, total bacterial load as assessed by 16S qPCR was also significantly higher in the LI than the SI (*p* < 0.0001), and also tended to be lower in SSD-fed mice than those fed chow or inulin (*p* = 0.09 for effect of diet; **Figure 8C**). Analysis of β-diversity again showed a significant interaction between diet and infection (**Figure 8D**; *p*<0.01 by PERMANOVA on Bray-Curtis distance metrics), however pairwise testing did not identify significant difference between groups following multiple comparison adjustment. Notably, the taxonomical composition of the LI closely reflected that observed after longer periods of diet consumption observed previously (Figures 4 and 5), suggesting rapid adaption of the GM to the tested diets (**Supplementary Figure 3**). The dominant genera in both the LI and SI were *Faecalibaculum, Bifidobacterium, Lactobacillus*, and *Muribaculaceae,* with the LI additionally harboring a number of genera such as *Bacteroides, Blautia, Alstipes* and *Roseburia* that were mostly absent in the SI (**Supplementary Figure 3**). Thus, whilst the exact mechanism(s) that lead to the altered pathogen loads remain to be revealed, our data show that the degree of diet-mediated susceptibility to infection correlates with total bacterial load at the infection site.

**Figure 8.**
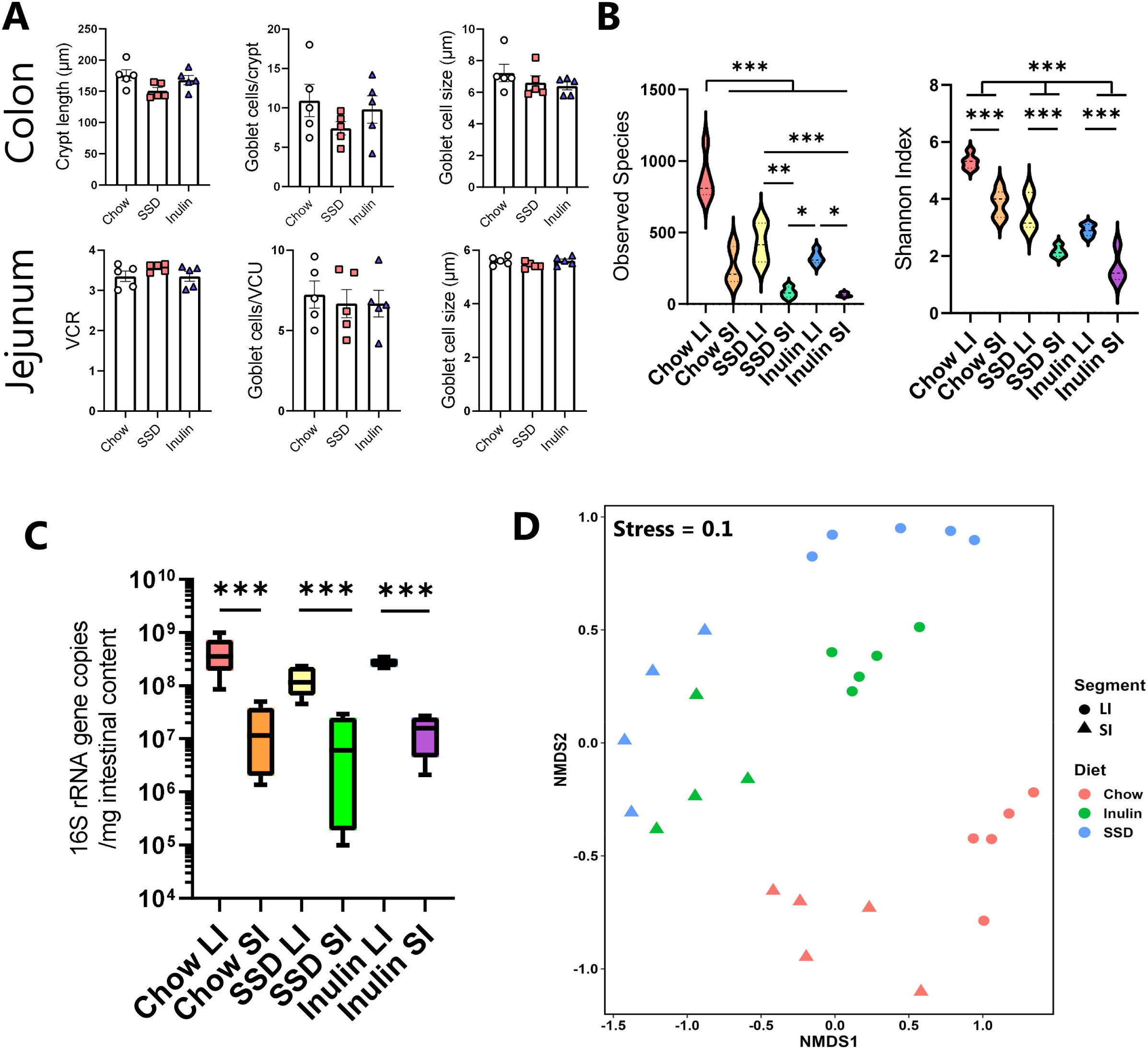
Impact of different diets on mucosal histology and microbiome composition in the small intestine and colon prior to infection. Mice were fed either chow, semi-synthetic diets alone (SSD), or SSD enriched with 10% inulin (Inulin) for 14 days. n=4-5 per group. **A)** Colonic crypt lengths, jejunum villous height to crypt depth ratio (VCR), and goblet cell numbers and size in the jejunum and colon. n=5 per group, shown are means ± S.E.M. **B)** α-diversity (Observed species and Shannon index), in the large (LI) and small intestine (SI) mice fed chow, SSD, or inulin. \*\*\**p*<0.005, \*\**p*<0.01, \**p*<0.05 by two-way ANOVA, with Tukey’s post-hoc testing where interaction term is significant. **C)** Non-metric multidimensional scaling (NMDS) plot showing β-diversity of different treatment groups. **D)** Total bacterial load in LI and SI in mice fed different diets, assessed by qPCR of 16S rRNA gene copy number. \*\*\**p*<0.005, by two-way ANOVA.

## Discussion

In the current work we present data which show that diet composition is a key factor determining the severity of enteric infections in the caecum and colon of mice. Our results build on a growing body of literature which suggest that intestinal infections may be highly sensitive to the levels of different dietary components.

The consumption of whole-grain foods and soluble fibers is generally considered to promote gut health through reducing inflammation and stimulating the growth of beneficial, metabolite-producing microbes, such as short chain fatty acids (SCFA)-producing gut bacteria (11). SCFA such as butyrate have been shown to have many beneficial effects, including providing a fuel source for oxidation in colonic epithelial cells, and suppressing the growth of pathogenic bacteria (41, 42). However, whilst this paradigm holds true in many pre-clinical models of chronic diseases, there is evidence that, in some contexts, putatively healthy dietary components may negatively impact the gut environment. For example, inclusion of refined inulin powder in SSD can exacerbate colitis and even lead to hepatic cancer in mice (43, 44), and we have found that the acute expulsion of a high dose of *T. muris* in C57BL/6 mice is impaired in mice fed inulin-enriched SSD (32). We have now extended this to show that whilst feeding rodent chow allows the establishment of a chronic *T. muris* trickle infection, as has previously been reported (28), feeding SSD results in substantially lower worm burdens and appears to promote stronger type-2 anti-helminth immunity. These results thus challenge the paradigm that whole-grain-containing diets with large amounts of unrefined plant material is protective against enteric pathogens. Interestingly, we also showed that this effect is also applicable to *C. rodentium.* Of note, several recent reports have demonstrated similar findings. An *et al.* also showed that colonization with *C. rodentium* was enhanced in mice fed either chow or inulin-enriched SSD (33), relative to SSD whilst Smith *et al.* have found that large amounts of resistant starch incorporated into SSD also promote higher *C. rodentium* burdens (45). Similarly, *Salmonella* burdens also appeared to be enhanced by chow in comparison to SSD (27).

The precise mechanisms whereby diet composition drives this enhanced susceptibility to infection remain yet unresolved, although our current data suggest that the mechanisms are different between *T. muris* and *C. rodentium*. An *et al.* proposed that feeding SSD to mice, which is low in fermentable fiber, results in a loss of commensal gut microbes and a nutrient-poor gut environment which does not support *C. rodentium* colonization (33). In contrast, the inclusion of fermentable fibers allows the production of microbial-derived metabolites, such as the SCFAs acetate and propionate. These metabolites, particularly acetate, can be directly utilized as energy sources by *C. rodentium* (33), providing a mechanistic basis for the increased *C. rodentium* colonization. However, An *et al.* also showed that whilst the initial establishment of *C. rodentium* is higher in mice fed chow in long-term experiments, these mice are capable of clearing the infection 2-3 weeks after colonization, whereas mice fed SSD retained a low-level chronic infection (33). This suggests that the presence of a rich and diverse microbiome allows early colonization with pathogenic bacteria but that the commensal microbes eventually outcompete and exclude the pathogen. In contrast, a nutrient-deprived gut environment shaped by SSD will result in lower initial establishment but those bacteria that do establish will persist over a longer period as there is less competition from commensals. This is in line with our experiments that show that increased *C. rodentium* establishment in inulin-fed mice is not accompanied by obvious defects in immunity as revealed by RNA-seq analysis, and is not reversed by treatment with IL-22, a known stimulator of anti-*Citrobacter* immunity (39). Interestingly, most of the top upregulated genes (e.g. *Cep85*) in inulin-fed mice infected with *C. rodentium* were related to basic cellular processes such as centromere function and other cell cycle activities. The reasons for this can only be speculated on, but one possibility is that the enhanced bacterial levels in these mice stimulate increased cellular turnover and repair mechanisms which necessitates the need for upregulation of pathways related to cell cycle processes. This proposed mechanism is clearly different to how chow and inulin enhance *T. muris* burdens. In this case, SSD results in rapid expulsion of the parasite whilst either chow or inulin consumption results in a long-term chronic infection, accompanied by a type-1 biased immune response which is known to be detrimental for helminth expulsion. In addition, we have previously shown that the increased numbers of *T. muris* worms in inulin-fed mice are not accompanied by increased SCFA production (32), suggesting that *T. muris* does not leverage inulin-induced SCFA production to promote its survival, in contrast to *C. rodentium*.

How diet composition drives this type-1 bias is an ongoing question. Immune cells in the gut are calibrated by continual exposure to both dietary compounds and GM-derived metabolites. The enrichment of these ligands through the consumption of more complex diets (as opposed to highly simple and refined SSD) may result in a tuning of host intestinal immune processes towards a type-1 response and away from the protective type-2 response. Evidence for switches in the type-1/type-2 axis in response to dietary cues have been reported. For example, differences in the levels of dietary vitamin A or the activation of the Aryl hydrocarbon receptor, a transcription factor highly sensitive to dietary compounds, can modulate innate lymphoid cell (ILC) balances to suppress ILC2 responses and render mice susceptible to helminths (46, 47). Of note, germ-free mice which are expected to have a very simple gut environment due to the absence of microbes to break down dietary fiber, have exaggerated type-2 immune responses, suggesting a continuum between the complexity of the gut metabolic environment and the degree of polarization towards a type-1 response. However, it should be noted that inulin can influence host immune function even in the absence of a gut microbiota (48). Further experiments will be necessary to unravel these complex interactions between diet, infections and the host immune system.

Our data also indicate that the enhancement of pathogen burdens by either chow or inulin-enriched SSD was more pronounced when infections were located in the large intestine rather than the small intestine. The reason(s) for this are not yet clear but may relate to the increased fermentative capacity of the caecum and colon relative to the small intestine, consequently exposing the pathogens to a more metabolically diverse environment. Interestingly, we observed that chow intake resulted in higher *G. muris* cyst output than SSD (regardless of inulin inclusion). The reasons for this, and why cyst output should be affected but not trophozoites numbers, are not clear and warrant further investigation. Potentially, a metabolic cue that stimulates the encysting of *G. muris* is suppressed when mice are fed SSD. Overall, there appears to be conserved effects of diet across multiple, evolutionary distinct pathogens but tissue niche may also be an important factor in diet-pathogen interactions. These conclusions are limited by the obvious fact that different pathogens are studied in the different gut compartments, and our data may simply reflect differences in pathogen life cycle and relationship with the host immune system, rather than an inherent effect of tissue predilection site.

In conclusion, we have shown that diet composition can affect enteric infection intensity in distinct murine infection models. The findings have clear relevance for the rational design of nutritional interventions with functional food components. Moreover, our results suggest that pharmacological or nutritional modulation of the gut environment may hold promise to develop novel therapeutic tools to limit infections and promote gut health.

## Acknowledgments

We are grateful to Pankaj Arora, Mette Schjelde, Penile Jensen, Stig Thamsborg, Emil Jakobsen, and Joshua Malsa for assistance. This work was supported by Independent Research Fund Denmark (Grants 7026-0094B and 2034-00245B), the Novo Nordisk Foundation (Grants 0052422 and NNF22OC0074714), and the A.P. Møller Foundation (Grant 20-L-0300).

## Methods

### Mice and pathogens

C57BL/6 mice aged 6-7 weeks were used for all experiments and were sourced from either Envigo for *T. muris*, *C. rodentium* or *H. polygyrus* experiments, or Charles River for *G. muris* experiments. Mice were fed either standard chow, or, where indicated, purified open-source SSD (13 kJ% fat (E15051); ssniff Spezialdilllten GmbH, Germany) or SSD with 10% long-chain inulin replacing corn starch, as reported previously (32). Mice were fed for two weeks before infections commenced. *T. muris*, *H. polygyrus, C. rodentium* and *G. muris* were propagated as described previously (30, 32, 49, 50). All infections were performed by oral gavage by dispensing a total volume of 0.2 ml per mouse. For *T. muris*, infections were performed by trickle-dosing with 20 eggs three times at seven-day intervals or, where mentioned, single doses of either 20 or 40 eggs. Infective doses for other pathogens were 10^9^ *C. rodentium* CFU, 200 *H. polygyrus* third-stage larvae (L_3_), or 10^4^ *G. muris* cysts. All animal experimentation was approved by relevant committees (License number 2020-15-0201-00465 for *T. muris*, *H. polygyrus* and *C. rodentium* infections and EC2019/077 for *G. muris* experiments). Mice weights were monitored weekly and welfare assessed daily by experienced personnel. We observed no significant effects of diets or infection on weight gain. Mice were sacrificed at the indicated times post-infection (p.i). *T. muris* and *H. polygyrus* burdens were determined with a dissecting microscope. Worm egg counts were assessed using a modified McMaster technique. *C. rodentium* loads in faecal samples were determined by serial dilution and plating overnight at 37°C on MacConkey agar. *G. muris* trophozoites were counted using a haemocytometer after incubating the small intestine in PBS on ice, with shaking.

### In vivo administration of cytokines

Where indicated, 5 µg of IL-25 (#587306; BioLegend), or sterile PBS as a vehicle control, was administered by intraperitoneal (i.p.) injection every other day between day 3 and day 11 following *T. muris* infection, and 5 µg of IL-22 (#576206; BioLegend), or PBS, was administered by i.p. injection on day 0, 2, and 4 following *C. rodentium* infection.

### Mesenteric lymph node collection and flow cytometry

Mesenteric lymph nodes (MLN) were dissected, passed through a 70 µm cell strainer and suspended in PBS supplemented with 2% foetal calf serum. Cells were washed, and surface stained with FITC-conjugated hamster anti-mouse TCRb (clone H57-597; BD Biosciences) and PerCP-Cy5.5-conjugated rat anti-CD4 (clone RM4-5; BD Biosciences), followed by intracellular staining with Alexa Fluor 647– conjugated mouse anti-mouse T-bet (clone 4B10; BD Biosciences), PE-conjugated rat anti-mouse GATA3 (clone TWAJ; Thermo Fisher Scientific), or FITC-conjugated rat anti-mouse Foxp3 (FJK-16s; Thermo Fisher Scientific). Cells were analyzed on a BD Accuri C6 flow cytometer (BD Biosciences), and data were analyzed using Accuri CFlow Plus software (Accuri Cytometers). Gating strategy is shown in Supplementary Figure 4.

### Histology

Colon or jejunum samples were fixed in 4% paraformaldehyde, before paraffin wax embedding, sectioning and Periodic acid-Schiff (PAS) staining. Goblet cells were determined in 5 colonic crypts or 5 jejunum villous-crypt units per mouse. Goblet cell diameters were measured for 10 randomly selected cells per mouse. Tissue sections were photographed using a Leica DFC480 camera and measurements performed using LAS v4.6 software (Leica, Switzerland).

### RNA extraction and qPCR

RNA was extracted from full-thickness caecum or colon tissues as previously described (30). cDNA was synthesized using Quantitect Reverse Transcription kits (Qiagen) and qPCR was carried out using the following program: 95°C for 2 min followed by 40 cycles of 15 s at 95°C and 20 s at 60°C. *Gapdh* was used as a reference gene for normalization and fold changes calculated using the ΔΔCT method. Primer sequences are listed in Table 1.

**Table 1.**
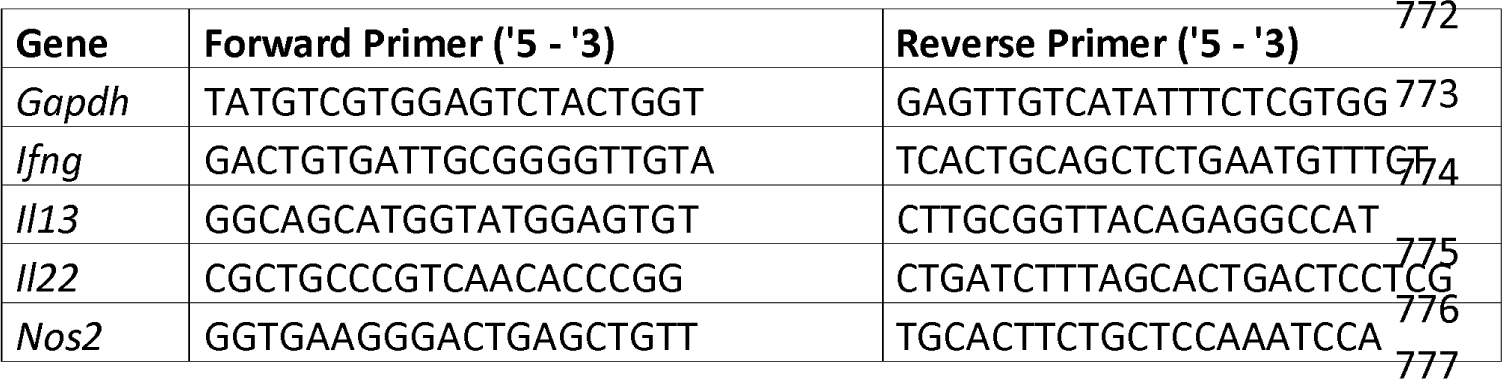
Primer sequences used for qPCR.

### RNA sequencing

RNA was used for 150bp paired-end Illumina NovaSeq6000 sequencing (Novogene, Cambridge, UK). Raw sequences were mapped to the mm10 genome assembly and read counts generated which were used to determine differentially expressed genes using DEseq2 (51). Pathway analysis was conducted using gene-set enrichment analysis (52). Volcano plots were constructed using R package ggplot2 (3.4.3) (53).

### 16S rRNA amplicon sequencing

Fresh faeces was collected after defecation from mice in the *T. muris* and *C. rodentium* studies. Alternatively, intestinal content from the small intestine (full length) or large intestine (caecum and proximal colon) was collected after 14 days diet adaption. Samples were immediately placed on ice before transfer to –80°C within one hour of collection. micDNA was extracted using FastDNA Spin kit for soil (MP Biomedicals, USA), including bead beating. 16S RNA Amplicons (regions 1-8, bV18-A) were prepared using the following primers: 8F - AGRGTTYGATYMTGGCTCAG, 1391R - GACGGGCGGTGWGTRCA. The PCR program was initial denaturation at 98 °C for 3 min, 25 cycles of amplification (98 °C for 30 s, 62 °C for 20 s, 72 °C for 2 min) and a final elongation at 72 °C for 5 min. Sequencing libraries were prepared from purified amplicons using the SQKLSK114 kit (OxfordNanopore Technologies, UK) according to manufacturer’s protocol with the following modifications: 500 ng total DNA was used as input, and CleanNGS SPRI beads (CleanNA, NL) were used for library cleanup steps. Libraries were loaded onto a MinION R10.4.1 flowcell and sequenced using MinKNOW 22.12.7 software (Oxford Nanopore Technologies, UK).

### Analysis of 16S rRNA amplicon data

The sequencing reads in the demultiplexed and basecalled fastq files were filtered for length (320 - 2000 bp fragments selected) and quality (phred score > 15) using a local implementation of filtlong (github.com//rrwick/Filtlong) v0.2.1 with the settings –min_length 320 – max_length 2000 – min_mean_q 97. The SILVA 16S/18S rRNA 138 SSURef NR99 full-length database in RESCRIPt format was downloaded from QIIME (54, 55). Potential generic place holders and dead-end taxonomic entries were cleared from the taxonomy flat file, i.e., entries containing uncultured, metagenome or unassigned, were replaced with a blank entry. The filtered reads were mapped to the SILVA 138.1 99 % NR database with minimap2 v2.24r-1122 using the -ax map-ont command (56) and downstream processing using samtools v1.14 (57). Mapping results were filtered such that query sequence length relative to alignment length deviated < 5 %. Noteworthy, low-abundant OTUs making up < 0.01 % of the total mapped reads within each sample were disregarded as a data denoising step. Further bioinformatic processing was done via RStudio IDE (2022.2.3.492) running R version 4.2.3 (20230315) and using the R packages: ampvis2 (2.7.27) (58), tidyverse (1.3.1), seqinr (4.2.16), ShortRead (1.54.0) and iNEXT (2.0.20) (59, 60). Analysis of β-diversity was carried out using NMIT and PERMANOVA (999 permutations) on Bray-Curtis dissimilarly metrics, followed by pairwise testing with Adonis using R package vegan (2.6.4) (61), applying Bonferroni corrections for adjusted *p* values.

### qPCR for total bacterial load

Total bacterial load was quantified using universal 16S rRNA probe and primers (F – TCCTACGGGAGGCAGCAGT, R – GGACTACCAGGGTATCTAATCCTGTT) as previously described (62), utilizing a linearized plasmid containing the PCR amplicon from *Escherichia coli* as standard.

### ELISA

Excretory/secretory antigens from *T. muris* and *H. polygyrus* were produced as previously described (32). *C. rodentium* antigens were produced by homogenization of cultured bacterial pellet in lysis buffer (CellLytic B, Sigma-Aldrich) followed by centrifugation and harvesting of the soluble protein antigens. Serum IgG1 and IgG2a specific for the different antigens was determined by ELISA as previously described (63).

### Statistical analysis

*P* values of <0.05 were considered significant. Assumptions of normality were checked through Shapiro-Wilk tests, or inspection of histogram plots and Kolmogorov-Smirnov tests of ANOVA residuals. Parametric data were analyzed using unpaired t-tests, or 2-way ANOVA and Tukey post-hoc testing, and presented as means ± S.E.M. Non-parametric data were log_10_ transformed prior to 2-way ANOVA, with the results presented as geometric means and 95% confidence intervals. Alternatively non-parametric data were analyzed using Mann-Whitney tests or Kruskal-Wallis and Dunn’s post hoc tests, and results presented as median values. Details of each experiment are given in the appropriate figure legends.

### Data availability

Raw RNA-sequence data is available at GEO under accession number GSE223377. 16S rRNA amplicon sequence data is available at SRA under BioProject Accession number PRJNA1016163.

