## Supplementary Figures for "Diet composition drives tissue-specific intensity of murine enteric infections"

### Supplementary Figure 1

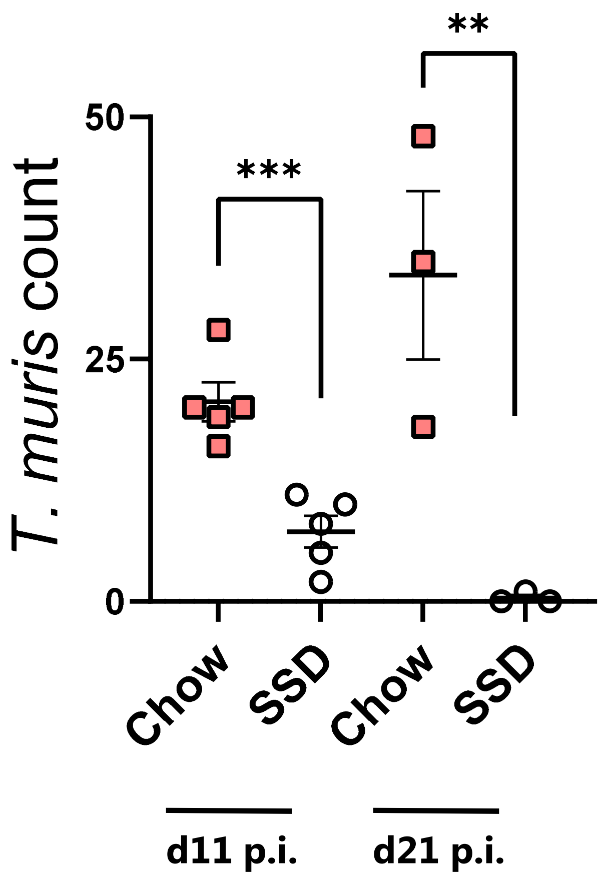


**Figure S1.** Mice were fed chow or SSD and infected with 40 *T. muris* eggs. Worm burdens were examined at day 11 p.i. (n=5 for each diet group) and day 21 p.i. (n=3 for each diet group). ***$p<0.005$, **$p<0.01$ by unpaired t-test.

### Supplementary Figure 2

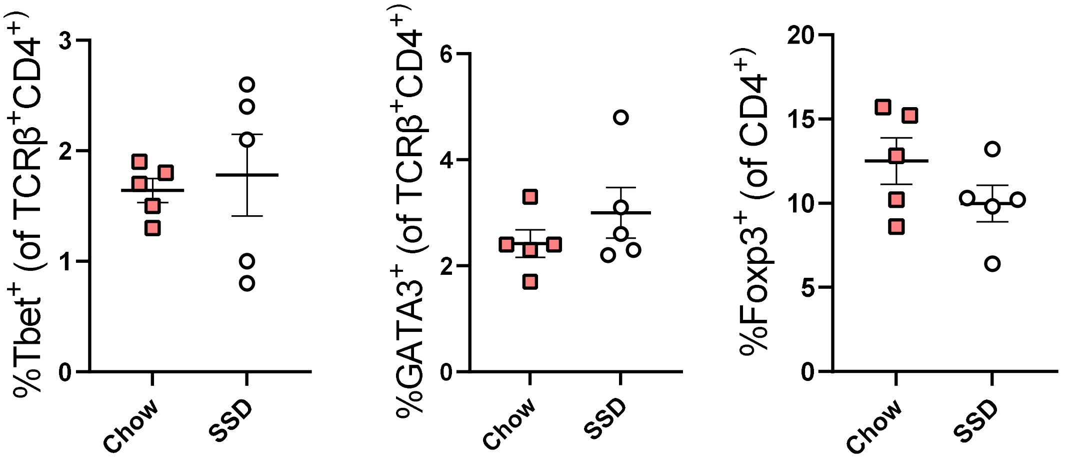


**Figure S2.** Mice were fed chow or SSD for 14 days (n=5 for each diet group). % of Th1 (Tbet$^+$), Th2 (GATA3$^+$) and Treg (Foxp3$^+$) cells in mesenteric lymph nodes were determined by flow cytometry.

### Supplementary Figure 3

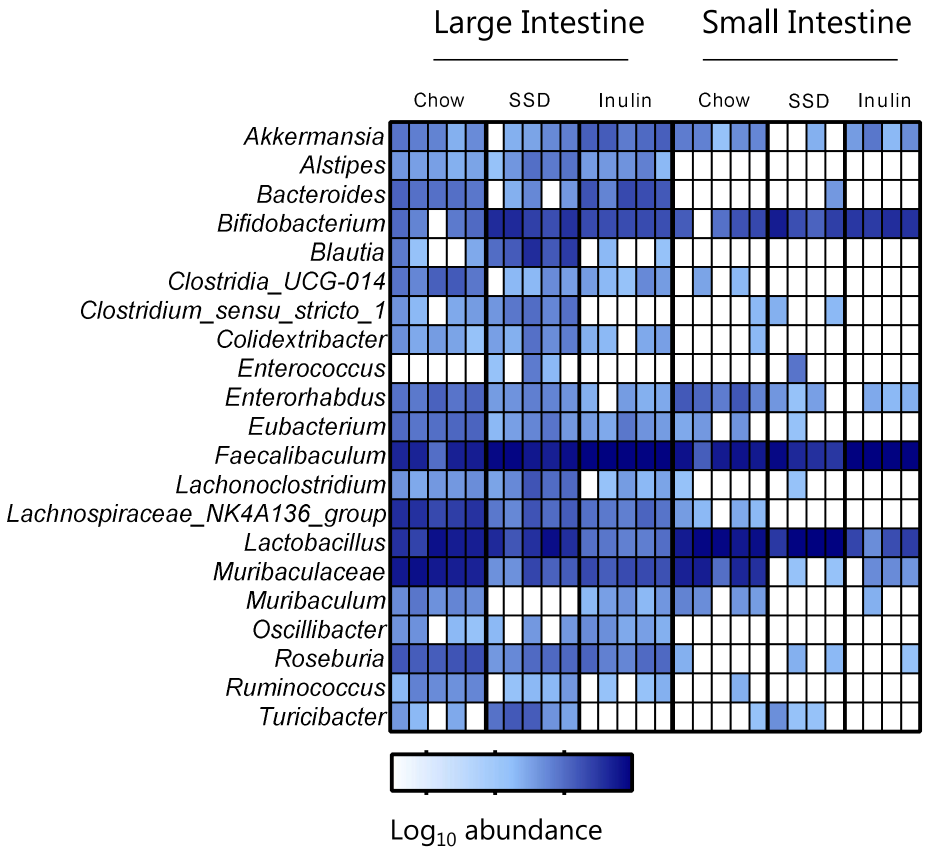


**Figure S3.** Mice were fed chow, SSD, or inulin-enriched SSD for 14 days (n=5 for each diet group). Gut microbiota composition was determined in the large intestine and small intestine, and presented as abundance at genus level.

### Supplementary Figure 4

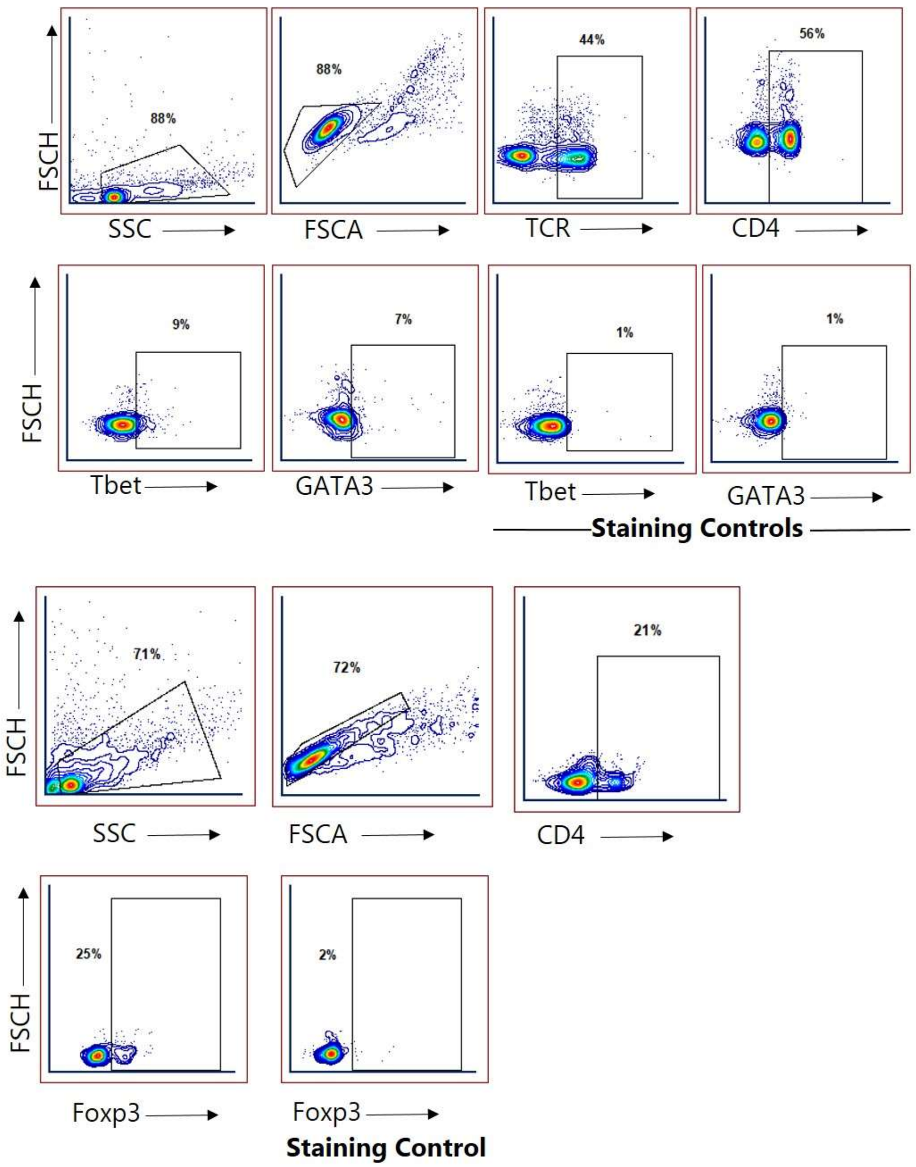


**Figure S4 –** Gating strategy for mesenteric lymph model cell flow cytometry analysis. Cells were gated on live cells and doublets excluded. For Tbet and GATA3 staining, cells were gated on TCR and CD4 expression, followed by intracellular staining for the transcription factors. For Foxp3 staining, cells were gated on CD4 followed by intracellular staining for the transcription factor.
